## Supplemental figure 1-11 for "Pu.1/Spi1 dosage controls the turnover and maintenance of microglia in zebrafish and mammals"

**This PDF file includes:**

Figs. S1 to S11

**A**

| | | Total number of fish | No. of GFP+ fish | Ratio |
|---|---|---|---|---|
| pre-selection at 3 dpf | | 147 | 43 | 29.3% |
| F0 screening efficiency | | 28 | 3 | 10.7% |
| F1 germline transmission ration | #1 | 77 | 28 | 36.4% |
| | #2 | 83 | 16 | 19.3% |
| | #3 | 47 | 2 | 4.3% |

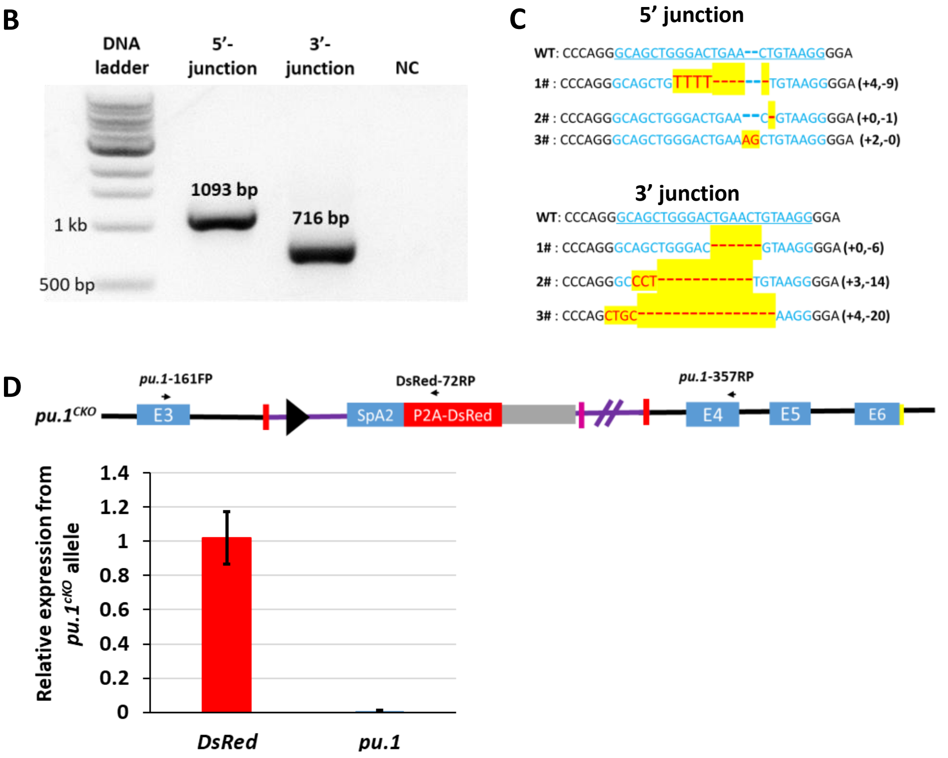


**Fig. S1. Characterization of *pu.1*$^{KI}$ and *pu.1*$^{CKO}$ alleles. (A)** Summary of the efficiency for the generation and screening of *pu.1*$^{KI}$. **(B)** Gel image shows the amplified 5'- and 3'-junction of the donor plasmid integration site in *pu.1*$^{KI}$ allele. **(C)** Summary of the genomic DNA sequences of the 5'- and 3'-junction in *pu.1*$^{KI}$ allele from different F0 founders. **(D)** Quantitative RT-PCR shows the relative expression of *DsRed* and *pu.1* from *pu.1*$^{CKO}$ allele.

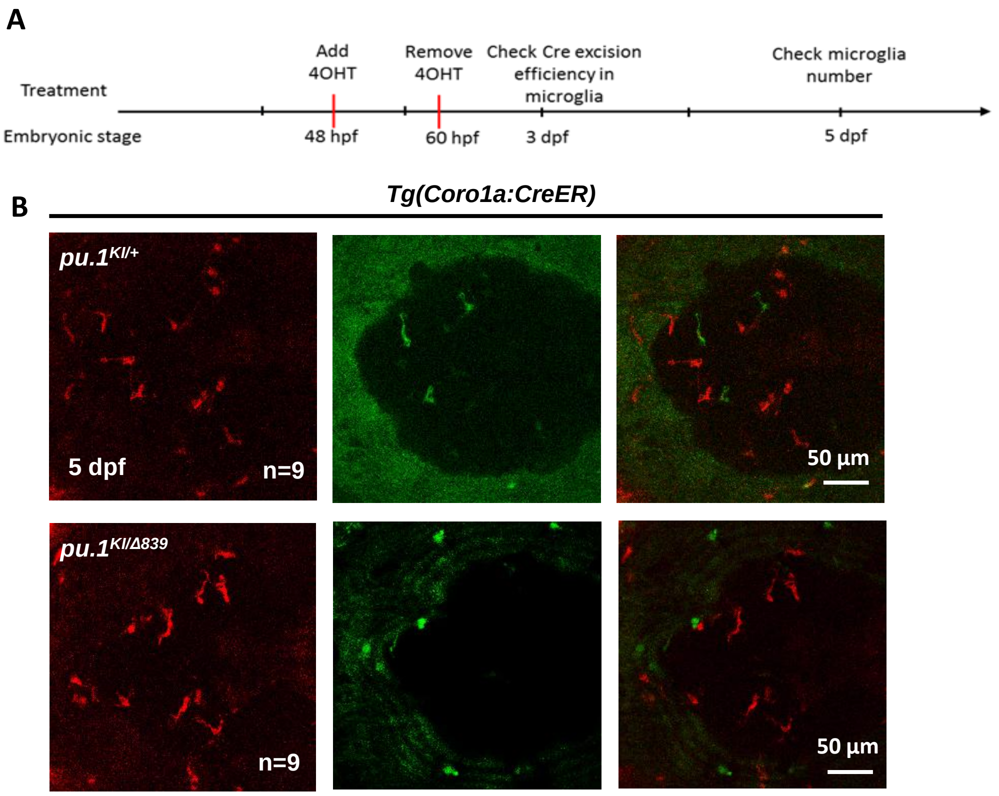


**Fig. S2. Conditional depletion of Pu.1 in embryonic microglia had no effect for their short-term survival.** **(A)** Schematics of 4-OHT treatment for *pu.1*$^{KI/WT}$ *Tg(coro1a:CreER)* and *pu.1*$^{KI/\Delta 839}$ *Tg(coro1a:CreER)* at embryonic stage. **(B)** Representative images of DsRed$^{+}$ microglia in *pu.1*$^{KI/WT}$ and *pu.1*$^{KI/\Delta 839}$ at 5 dpf.

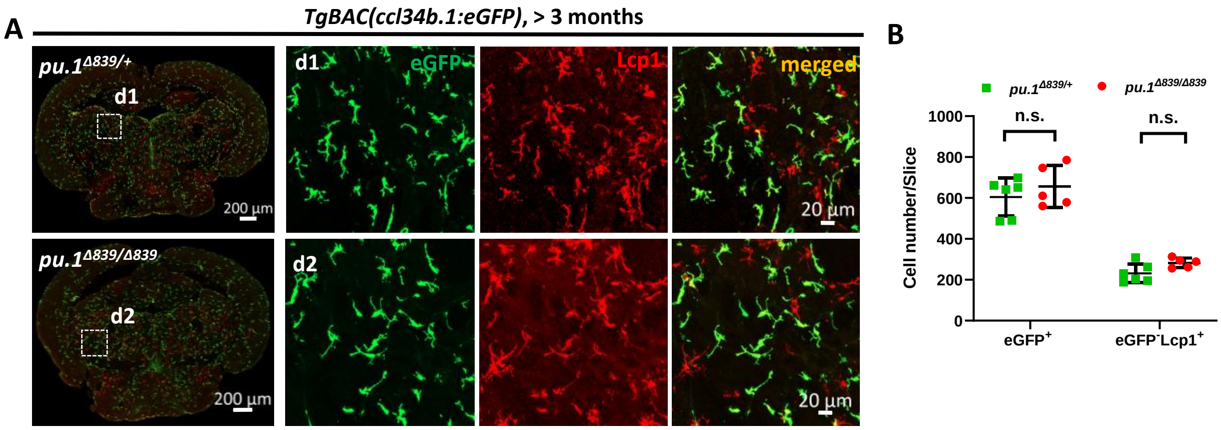


**Fig. S3. Microglia number is not affected in *pu.1*$^{\Delta 839}$ null mutants. (A)** Representative images show the co-staining of eGFP and Lcp1 antibodies on the midbrain cross section of adult *pu.1*$^{\Delta 839/+}$*;TgBAC(ccl34b.1:eGFP)* and *pu.1*$^{\Delta 839/\Delta 839}$*;TgBAC(ccl34b.1:eGFP)* fish. eGFP$^{+}$ cells represent microglia, whereas eGFP$^{-}$Lcp1$^{+}$ cells are DCs. **(B)** Quantification of the number of eGFP$^{+}$ microglia and eGFP$^{-}$Lcp1$^{+}$ DCs on the midbrain cross section of adult *pu.1*$^{\Delta 839/+}$*;TgBAC(ccl34b.1:eGFP)* (n=6) and *pu.1*$^{\Delta 839/\Delta 839}$*;TgBAC(ccl34b.1:eGFP)* (n=5) fish. Student t test. n.s. = not significant, p>0.05.

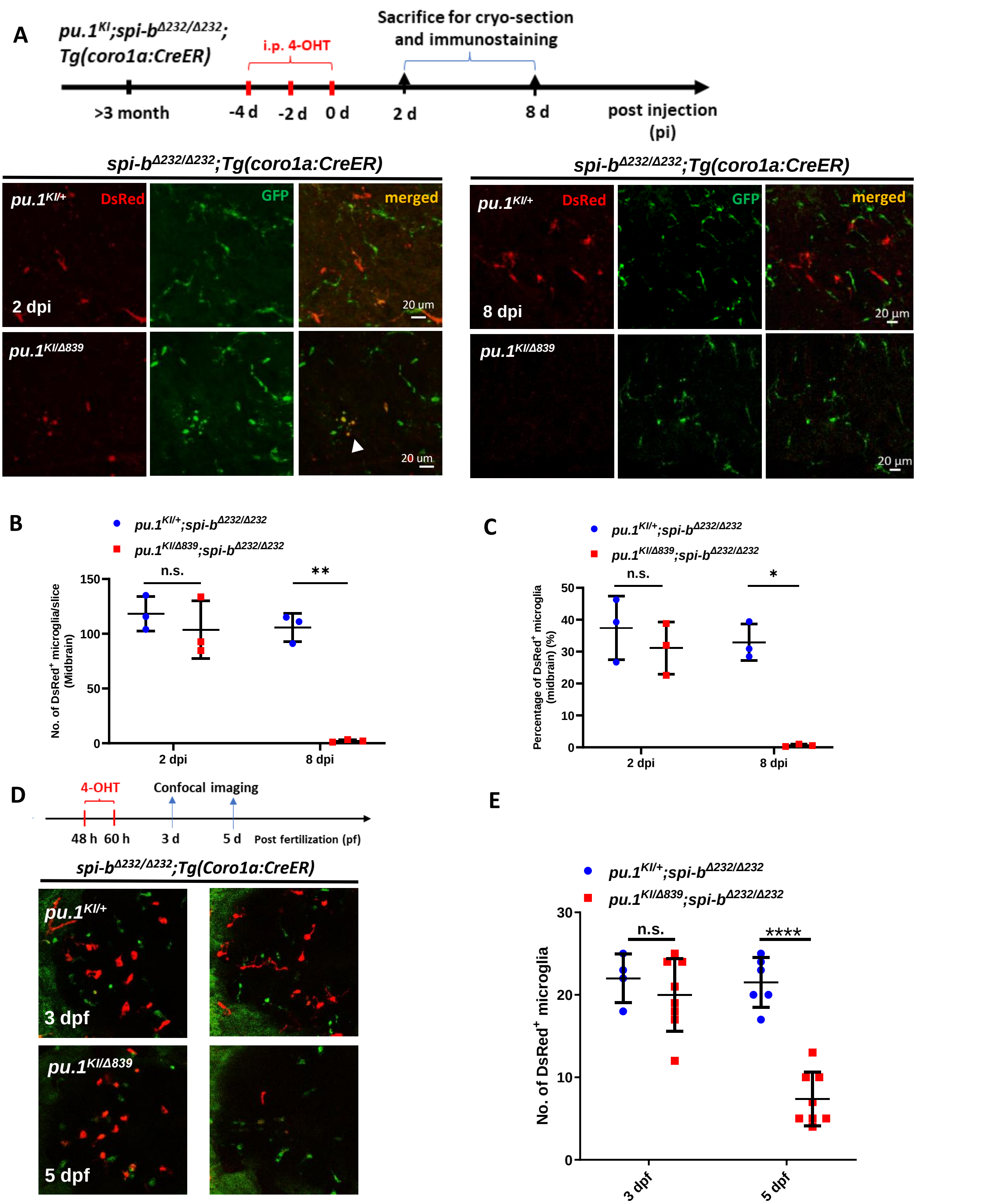


**Fig. S4. Simultaneous inactivation of Pu.1 and Spi-b lead to rapid elimination of microglia in zebrafish. (A)** The experimental setup for *pu.1* conditional knockout in adult $spi\text{-}b^{\Delta232/\Delta232}$ mutants and the representative images of the midbrain cross section of adult *$pu.1^{KI/+}$;$spi\text{-}b^{\Delta232/\Delta232}$;Tg(coro1a:CreER)* and *$pu.1^{KI/\Delta839}$;$spi\text{-}b^{\Delta232/\Delta232}$;Tg(coro1a:CreER)* fish at 2 dpi and 8 dpi. The white arrow indicates microglia with blebbing morphology. **(B-C)** Quantification of the number (B) and percentage (C) of DsRed$^{+}$ microglia in adult *$pu.1^{KI/+}$;$spi\text{-}b^{\Delta232/\Delta232}$;Tg(coro1a:CreER)* and *$pu.1^{KI/\Delta839}$;$spi\text{-}b^{\Delta232/\Delta232}$;Tg(coro1a:CreER)* fish at 2 and 8 dpi (n=3 for all groups). **(D)** The experimental setup for *pu.1* conditional knockout in $spi\text{-}b^{\Delta232/\Delta232}$ mutant embryos and the representative images show the DsRed+ microglia in *$pu.1^{KI/+}$;$spi\text{-}b^{\Delta232/\Delta232}$;Tg(coro1a:CreER)* and *$pu.1^{KI/\Delta839}$;$spi\text{-}b^{\Delta232/\Delta232}$;Tg(coro1a:CreER)* embryos at 3 and 5 dpf. **(E)** Quantification of the number of DsRed+ microglia in *$pu.1^{KI/+}$;$spi\text{-}b^{\Delta232/\Delta232}$;Tg(coro1a:CreER)* and *$pu.1^{KI/\Delta839}$;$spi\text{-}b^{\Delta232/\Delta232}$;Tg(coro1a:CreER)* embryos at 3 and 5 dpf after 4-OHT treatment from 48 to 60 hpf. (*$pu.1^{KI/+}$;$spi\text{-}b^{\Delta232/\Delta232}$;Tg(coro1a:CreER)* n=4 and 6 for 3 and 5 dpf respectively. *$pu.1^{KI/\Delta839}$;$spi\text{-}b^{\Delta232/\Delta232}$;Tg(coro1a:CreER)* n=8 for both 3 and 5 dpf.) Student t test. n.s. = not significant, $p>0.05$;***$p<0.001$;****$p<0.0001$

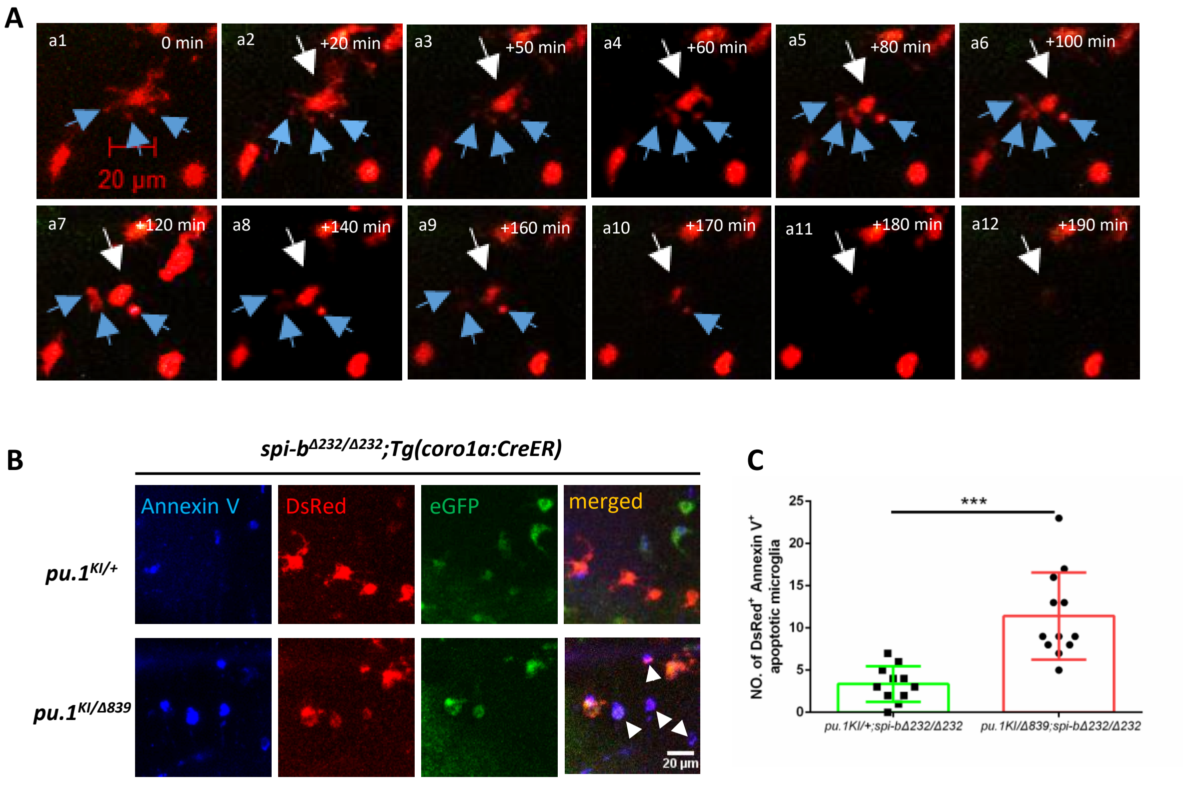


**Fig. S5. Pu.1/Spi-b-deficient microglia undergo apoptosis in zebrafish. (A)** Time-lapse live imaging shows the blebbing and fragmentation of DsRed$^{+}$ microglia between 3 dpf and 5 dpf in *pu.1*$^{KI/\Delta 839}$*;spi-b*$^{\Delta 232/\Delta 232}$*;Tg(coro1a:CreER)* embryos treated with 4-OHT from 48 hpf to 60 hpf. The blue arrows indicate the formation of apoptotic cell bodies. **(B)** Fluorescent live-imaging of Annexin V, DsRed and eGFP signals in 4-dpf *pu.1*$^{KI/+}$*;spi-b*$^{\Delta 232/\Delta 232}$*;Tg(coro1a:CreER) and pu.1*$^{KI/\Delta 839}$*;spi-b*$^{\Delta 232/\Delta 232}$*;Tg(coro1a:CreER)* embryos treated with 4-OHT from 48 hpf to 60 hpf and subjected to brain injection of Annexin V-647. **(C)** Quantification of DsRed$^{+}$Annexin$^{+}$ microglia in B. (*pu.1*$^{KI/+}$*;spi-b*$^{\Delta 232/\Delta 232}$*;Tg(coro1a:CreER)* n=11, *pu.1*$^{KI/\Delta 839}$*;spi-b*$^{\Delta 232/\Delta 232}$*;Tg(coro1a:CreER)* n=12) student t test. ***$p<0.001$.

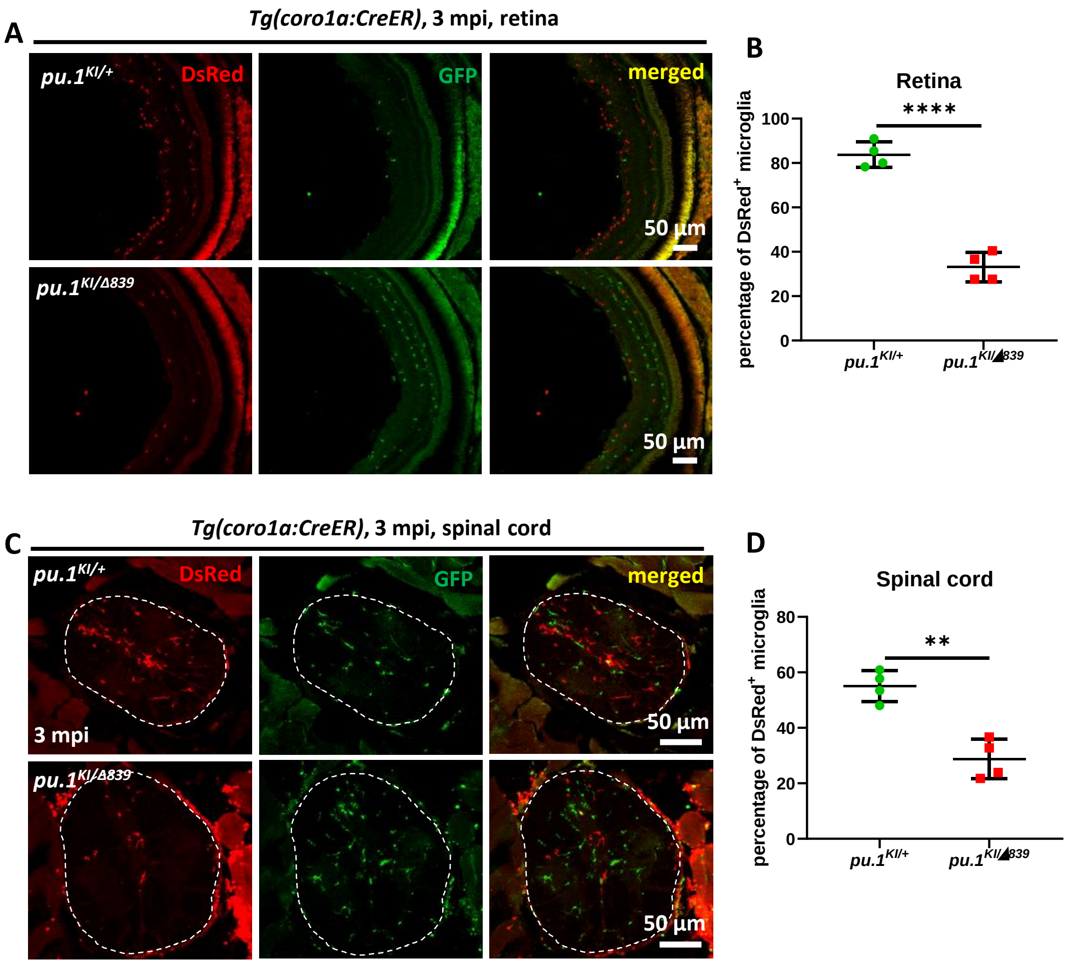


**Fig. S6. Conditional inactivation of Pu.1 leads to chronic elimination of microglia in the spinal cord and retina of adult zebrafish. (A)** Representative images of the retina cross section of adult *pu.1$^{KI/+}$;Tg(coro1a:CreER)* and *pu.1$^{KI/\Delta 839}$;Tg(coro1a:CreER)* fish at 3 mpi. **(B)** Quantification of the percentage of DsRed$^{+}$ microglia in the retina of adult *pu.1$^{KI/+}$;Tg(coro1a:CreER)* (n=4) and *pu.1$^{KI/\Delta 839}$;Tg(coro1a:CreER)* (n=4) fish at 3 mpi. **(C)** Representative images of the spinal cord cross section of adult *pu.1$^{KI/+}$;Tg(coro1a:CreER)* and *pu.1$^{KI/\Delta 839}$;Tg(coro1a:CreER)* fish at 3 mpi. **(D)** Quantification of the percentage of DsRed$^{+}$ microglia in the spinal cord of adult *pu.1$^{KI/+}$;Tg(coro1a:CreER)* (n=4) and *pu.1$^{KI/\Delta 839}$;Tg(coro1a:CreER)* (n=4) fish at 3 mpi. Student t test. **p<0.01; ****p<0.0001.

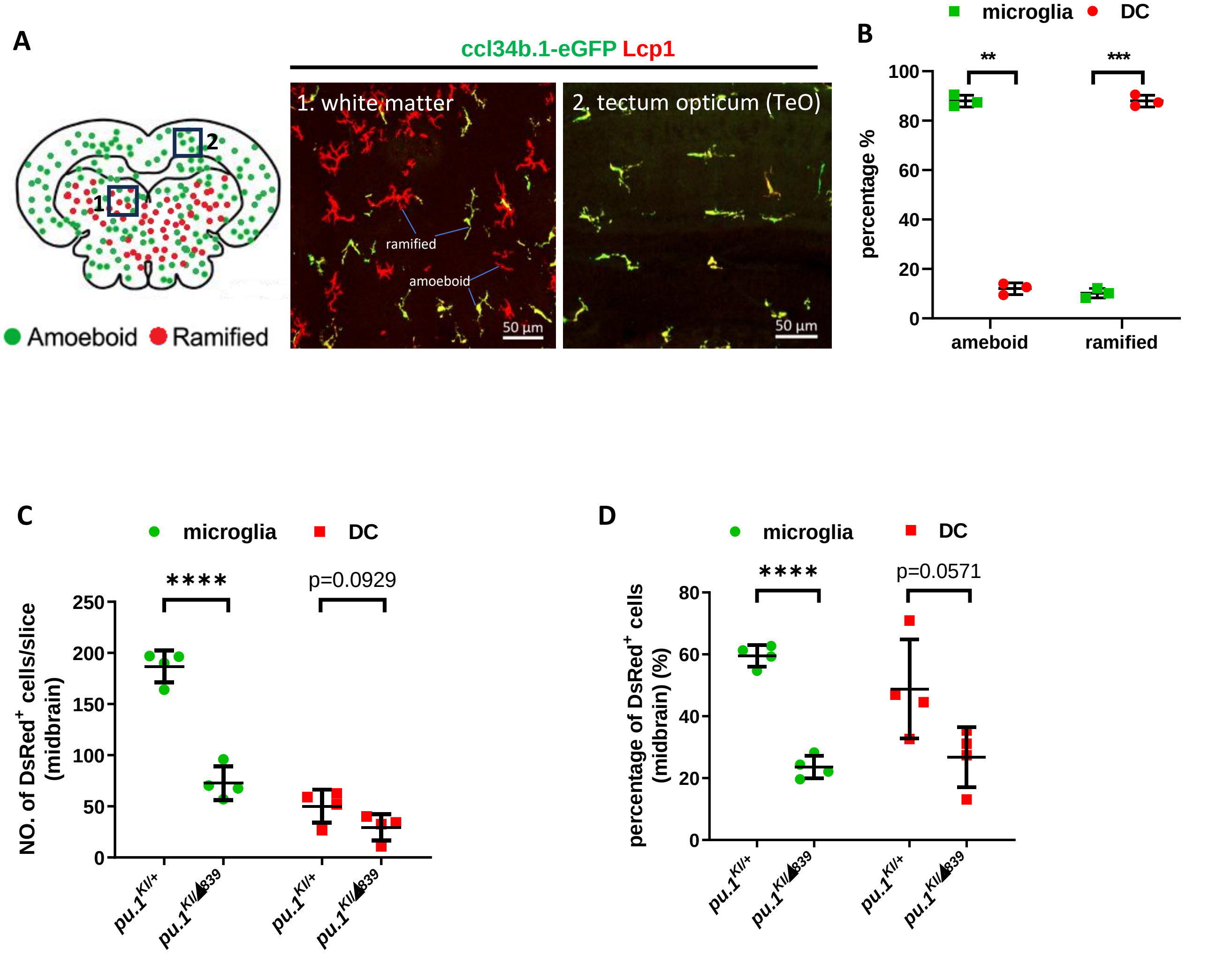


**Fig. S7. Conditional inactivation of Pu.1 leads to chronic elimination of microglia in the brain of adult zebrafish. (A)** Representative images showing different morphology of microglia (ccl34b.1-eGFP$^{+}$Lcp1$^{+}$) and DCs (ccl34b.1-eGFP$^{-}$Lcp1$^{+}$) in two midbrain regions of TgBAC*(ccl34b.1:eGFP)* fish. The amoeboid and ramified cells were indicated accordingly. **(B)** Quantification of the proportion of microglia (ccl34b.1-eGFP$^{+}$Lcp1$^{+}$) (n=3) and DCs (n=3) (ccl34b.1-eGFP$^{-}$Lcp1$^{+}$) in total amoeboid and ramified Lcp1$^{+}$ cells in the midbrain of TgBAC*(ccl34b.1:eGFP)* fish. **(C)** Quantification of the number of DsRed$^{+}$ microglia and DCs on the midbrain cross section of *pu.1*$^{KI/+}$*;Tg(coro1a:CreER)* (n=4) and *pu.1*$^{KI/\Delta 839}$*;Tg(coro1a:CreER)* (n=4) fish at 3 mpi by amoeboid and ramified morphologies. **(D)** Quantification of the proportion of DsRed$^{+}$ microglia and DCs on the midbrain cross section of *pu.1*$^{KI/+}$*;Tg(coro1a:CreER)* (n=4) and *pu.1*$^{KI/\Delta 839}$*;Tg(coro1a:CreER)* (n=4) fish at 3 mpi by amoeboid and ramified morphologies.

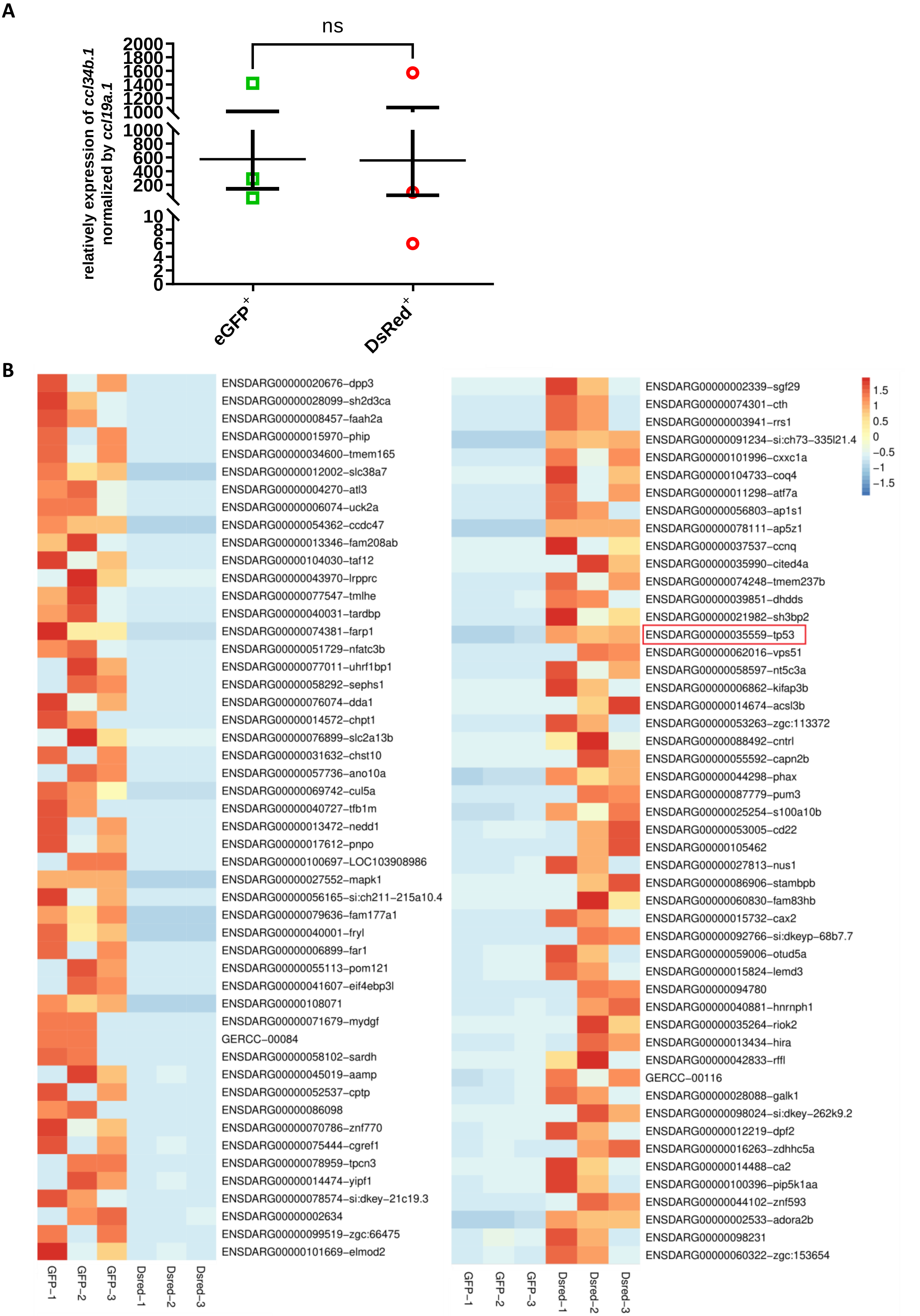


**Fig. S8. RNA-seq analysis of pu.1-deficient microglia. (A)** Relative expression of *ccl34b.1* normalized by *ccl19a.1* in the picked samples for RNA-seq analysis. **(B)** TPM heatmap of the top 50 differentially expressed genes in eGFP$^+$ and Dsred$^+$ microglia. Values in the heatmap are centered and scaled in the row direction. The red color represents high expression while the blue color represents low expression.

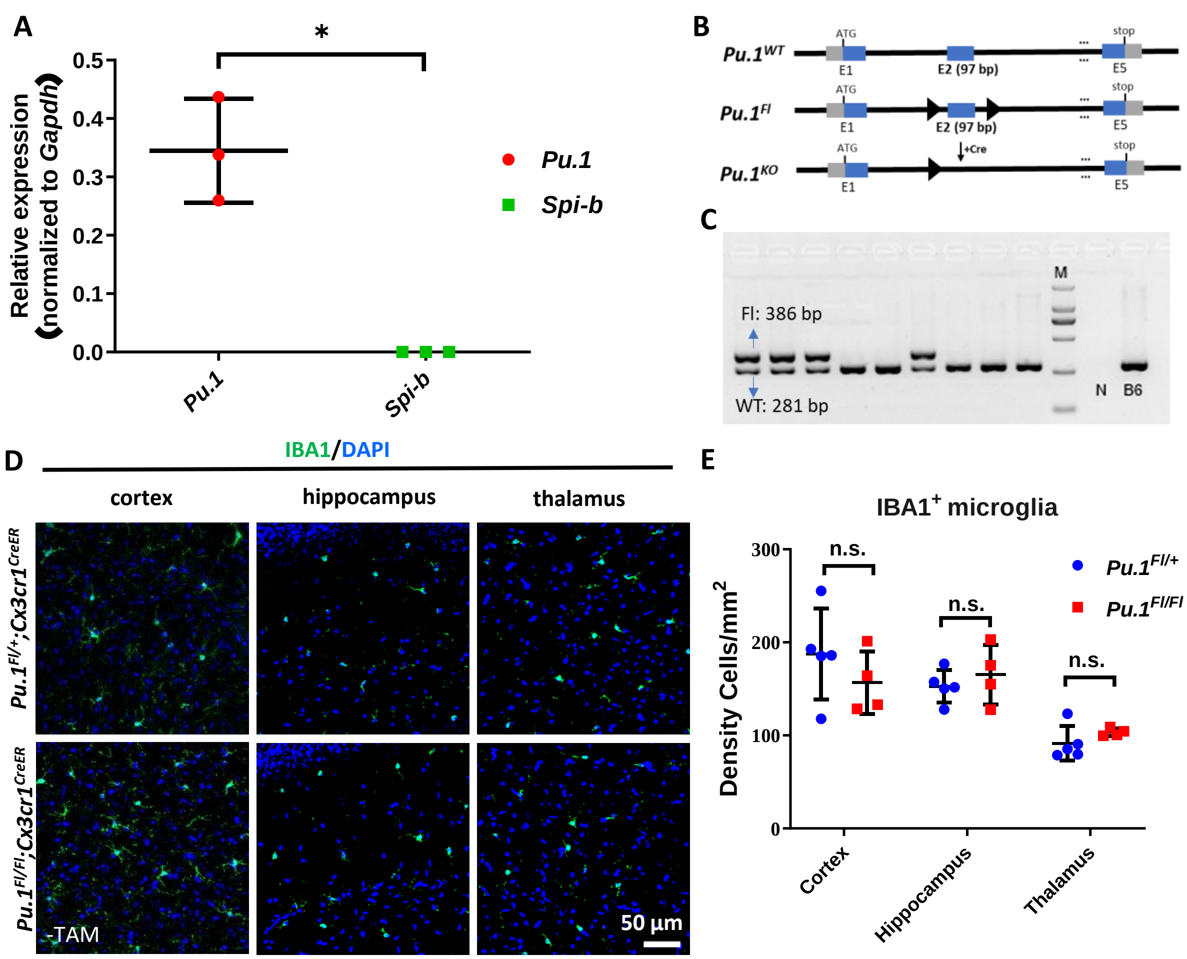


**Fig. S9. Adult microglia are not affected in *Pu.1$^{Fl/Fl}$;Cx3cr1$^{CreER}$* mice. (A)** Quantitative RT-PCR result shows the relative expression of *Pu.1* (n=3) and *Spi-b* (n=3) (normalized to *Gapdh* expression) in sorted microglia from adult mouse brain. **(B)** Schematic diagram shows the genomic loci of *Pu.1$^{WT}$*, *Pu.1$^{Fl}$* and *Pu1$^{KO}$* alleles respectively. **(C)** Gel image shows the genotyping result of *Pu.1$^{WT}$* and *Pu.1$^{Fl}$* alleles. **(D)** Representative images of IBA1 and DAPI co-staining in the cortex, hippocampus and thalamus regions of adult *Pu.1$^{Fl+/}$;Cx3cr1$^{CreER}$* and *Pu.1$^{Fl/Fl}$;Cx3cr1$^{CreER}$*mice without TAM injection. **(E)** Quantification of the Density (number) of IBA1$^{+}$ microglia in the cortex, hippocampus and thalamus regions of adult *Pu.1$^{Fl/+}$;Cx3cr1$^{CreER}$* (n=5) and *Pu.1$^{Fl/Fl}$;Cx3cr1$^{CreER}$* (n=4) mice. Student t test. n.s. = not significant, $p>0.05$. $*p<0.05$.

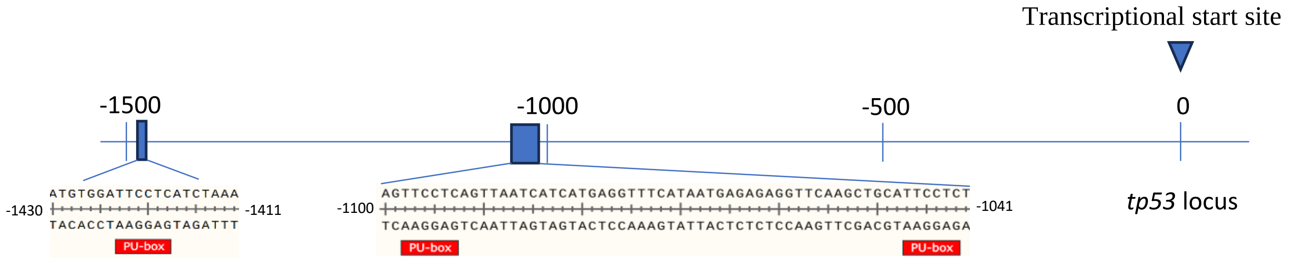


**Fig. S10. In-silico analysis of Pu.1 binding sites on the promoter region of *tp53*.** Three PU.1 binding sites (GAGGAA) locating on the antisense strand from position -1423 to -1418, -1098 to -1093 and -1047 to -1042 relative to the transcriptional start site of *tp53* were indicated by red box.

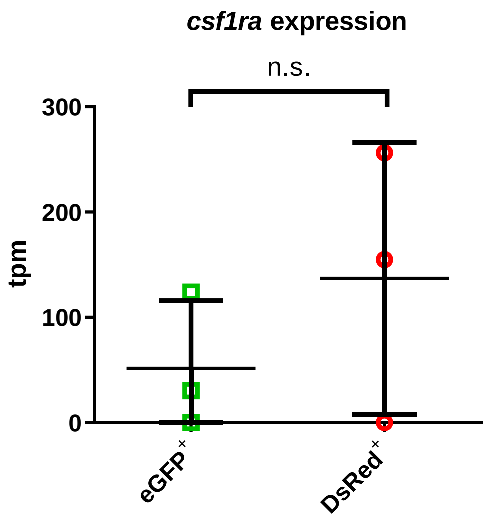


**Fig. S11 *csf1ra* expression does not decrease after conditional inactivation of Pu.1.** Relative expression of *csf1ra* in eGFP+ (n=3) and DsRed+ (n=3) microglia at 3 mpi by transcripts per million (TPM).
